# Massively parallel screening of TIR-derived peptides reveals vast TLR-targeting immunomodulatory peptides

**DOI:** 10.1101/2024.05.21.595261

**Authors:** Yun Lim, Tae Kyeom Kang, Meong Il Kim, Dohyeon Kim, Ji-Yul Kim, Sang Hoon Jung, Keunwan Park, Wook-Bin Lee, Moon-Hyeong Seo

**Author notes:** These authors contributed equally (Yun Lim, Tae Kyeom Kang). Corresponding authors: Moon-Hyeong Seo:; Wook-Bin Lee:; Keunwan Park.

## Abstract

Toll-like receptors (TLRs) are critical regulators of the immune system, and altered TLR responses lead to a variety of inflammatory diseases. Interference of intracellular TLR signaling, which is mediated by multiple Toll/interleukin-1 receptor (TIR) domains on all TLRs and TLR adapters, is an effective therapeutic strategy against immune dysregulation. Peptides that inhibit TIR-TIR interactions by fragmenting interface residues have potential as therapeutic decoys. However, a systematic method for discovering TIR-targeting moieties has been elusive, limiting exploration of the vast unsequenced space of the TIR domain family. Here, we developed a comprehensive parallel screening method to uncover novel TIR-binding peptides derived from previously unexplored surfaces on a wide range of TIR domains. We constructed a large peptide library, named TIR surfacesome, by tiling surface sequences of the large TIR domain family and screening against MAL^TIR^ and MyD88^TIR^, TIRs of two major TLR adaptor proteins, resulting in the discovery of hundreds of TIR-binding peptides. The selected peptides inhibited TLR signaling, demonstrated anti-inflammatory effects in macrophages and therapeutic potential in mouse inflammatory models. This approach may facilitate the development of TLR-targeted therapeutics.

## Introduction

Toll-like receptors (TLRs) are a family of pattern recognition receptors that play a crucial role in both innate and adaptive immunity by recognizing a wide range of external pathogens or endogenous damage-associated molecules^1,2^. The pathological consequences of dysregulated TLRs have been highlighted in chronic immune-related disorders, such as autoimmune diseases, infections, and neuroinflammation^1,3–5^. Owing to their critical role in inflammation, modulation of uncontrolled TLR signaling using TLR-targeting agents, including small molecules and peptides, can help overcome the progression of various inflammatory conditions and support immune homeostasis^6–13^. Following TLR activation, intracellular signaling occurs via oligomerization of cytosolic Toll/interleukin-1 receptor (TIR) domains present in TLRs and adapter proteins. TIR-mediated interactions induce the higher-order assembly of TLR signalosomes, such as myddosome and triffosome, to activate key transcription factors in inflammation, including nuclear factor-κB (NF-κB), activator protein 1 (AP-1), and interferon regulatory factors (IRFs). Therefore, blocking intracellular TIR-TIR interactions is a valid approach to target TLR signaling^11^.

TIR domains are an essential and widespread component of innate immunity across the evolutionary tree including animals, plants, bacteria, and archaea^2,14^. Structurally, the consensus TIR domain consists of a central five parallel β-sheet and surrounding five α-helices with surface-exposed connecting loops. Structural analysis of TIR domains has revealed that four key regions of the TIR surface—site 1 (AB and BB loops, and β-strand B), site 2 (α-helices B and C), site 3 (DD loop, and α-helix D), and site 4 (β-strand E, EE loop, and α-helix E)— mediate homo-or heterotypic interactions that lead to formation of TLR signaling complexes^11^. To elucidate the interfaces, researchers extensively screened TIR-derived peptide fragments from the surface-exposed regions in the human TIR domains^15–21^, which led to the discovery of potential TLR-inhibitory peptides that competitively block the TIR-TIR interfaces. These TIR-targeting peptides, when fused to a cell-permeable peptide, demonstrated *in vitro* and *in vivo* immunomodulatory activity in several inflammatory diseases by interfering with intracellular TIR-TIR interactions^11^. In addition, fragmented peptides from viral or bacterial proteins that bind human TIR domains to evade host immunity have been reported to inhibit TLR3, 4, 7, or 9^11^. Nevertheless, the investigation of potential cross-species TIR-TIR interactions has received limited attention in the field of molecular evolution and infectious disease^2,14^. Considering the large number of organisms with TIR homologous domains^14^ or TIR-binding proteins, along with the multivalent/multispecific interactions among TIRs, vast stretches of the potential TIR interfaces are yet to be sequenced. However, all TIR-targeting peptides reported to date have been selected using low-throughput approaches with only a few synthetic peptides derived from a small fraction of the TIR-binding interface. Additionally, the low sequence similarities of the reported TIR-targeting peptides have made it challenging to either predict or design new peptides that take advantage of the wide variety of TIR domains found in nature.

Massively parallel screening of short peptides is an emerging strategy for identifying motifs that mediate protein-protein interactions (PPIs)^22–27^, regulate target proteins^28^, or modulate PPIs in various disease models^29–31^. Recent advances in DNA synthesis and large-scale sequencing have facilitated systematic and comprehensive discovery of peptide binders. Fragment-based high-throughput screening has the potential to effectively uncover previously unknown cross-species TIR-TIR interactions and identify TIR-targeting peptides. In this study, to systematically identify immunomodulatory peptides targeting TLR signaling, we constructed TIR surfacesome (T-Surf), a massive TIR-derived peptide library, by tiling surface regions of the TIR domains across various species. The parallel screening of more than 190,000 peptides allowed for extensive exploration of TIR-binding sequences, yielding a large dataset of TIR-binding motifs. Several synthetic peptides that bind to MAL^TIR^ or MyD88^TIR^ from the peptide-phage display showed significant anti-inflammatory activity by inhibiting TLR pathways in macrophages. The therapeutic potential of the candidate peptides was further validated in mouse models including lipopolysaccharide (LPS)-induced sepsis and age-related macular degeneration. Thus, the peptides represent valuable lead compounds for both diseases. The screening technique developed in this study may facilitate peptide drug discovery for a broader range of inflammatory diseases by immunomodulation of TLR signaling.

## Results

### T-Surf library design and construction

We designed a library, ranging from 16-to 21-amino acid-long peptides, representing surface-exposed residues of TIR homology domains by fragmenting the 11 regions of TIRs defined in a previous study^11^ (**Fig. 1a** and **Supplementary** Fig. 1). First, a total of 779,406 fragmented unique peptides from 13,644 TIR domains were collected based on the TIR domain family sequences archived in Pfam PF01582. Next, peptides representing clusters of sequences differing by two or fewer amino acids were extracted. Tiled sequences from 14 human TIR domains and 5 putative TIR-binding domains (**Supplementary Table 1**) were additionally included in the final list to determine whether the previously reported human TIR-derived decoy peptides outcompeted the others from non-human TIRs. The final designed T-Surf library consisted of 190,945 peptides (**Dataset 1**) derived from 13,603 TIR domains (99.7% of the 13,644 domains in Pfam PF01582) (**Supplementary Table 2**). Including the sequences omitted by clustering, the T-Surf library covered 70.6% of the total TIR domain family sequences (**Supplementary Table 2**).

**Figure 1.**
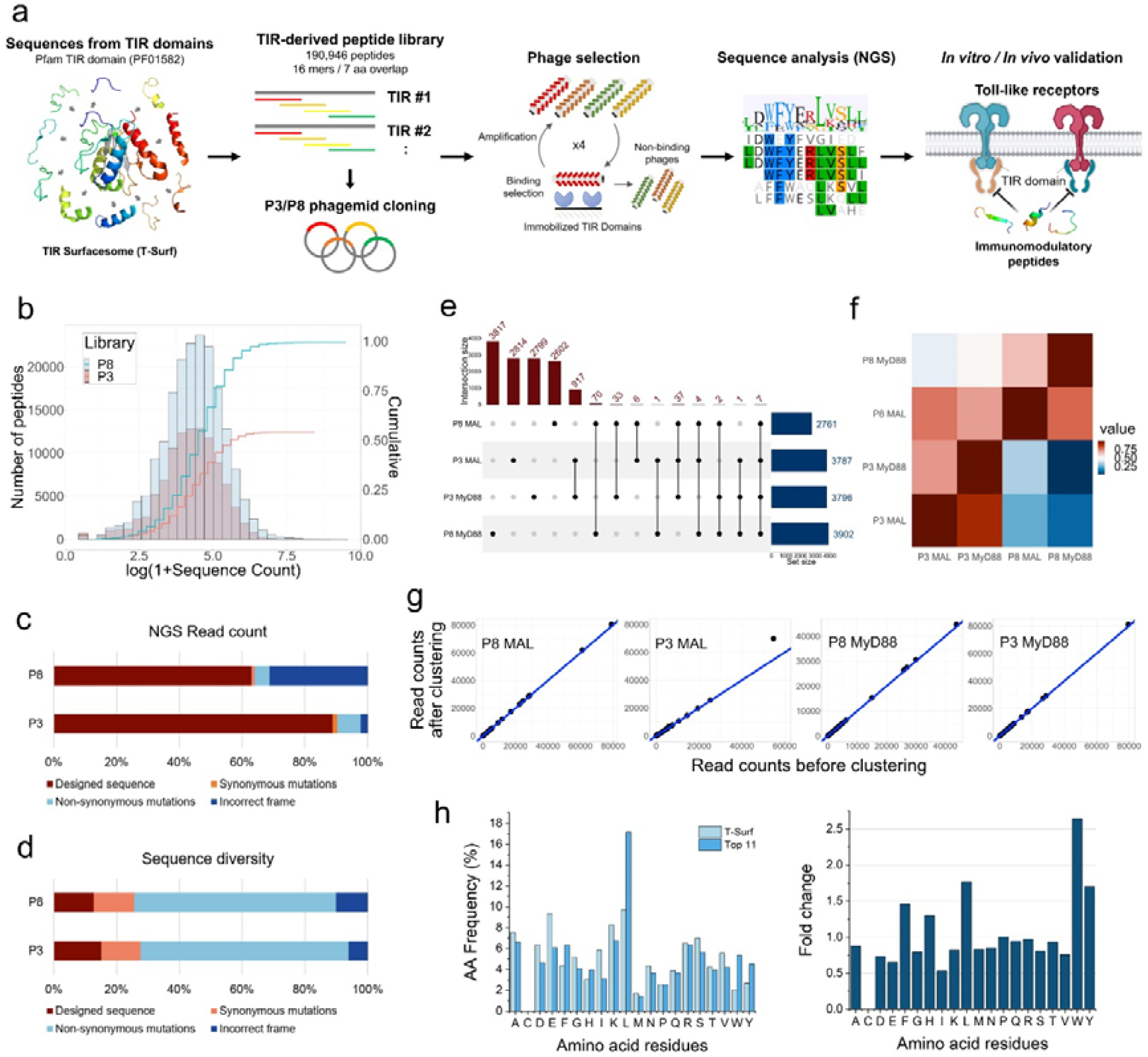
Design, construction, and selection of the TIR Surfacesome peptide library. (a) Schematic of the overall experiments. Fragmented peptide sequences exposed on the surface of the TIR domain family were collected to construct the TIR Surfacesome (T-Surf) library. The library was used to select TIR-binding peptides that modulate TLR signaling pathways by interfering with TIR-TIR interactions. After NGS analysis of the enriched peptides, their immunomodulatory activity was validated *in vitro* and *in vivo* models. (b) NGS analysis of the T-Surf naïve library. The count distribution of the T-Surf libraries and the coverage of the designed peptides as a function of the read counts are shown. (c) The ratio of NGS read counts of the libraries was analyzed by grouping the population into designed peptides, peptides containing silent or non-silent mutations, or incorrect frames containing stop codon or indel mutations. (d) The sequence diversity of the four classes was analyzed. (e) After four rounds of selection, the enriched peptide sequences were compared among the four pools. The number of sequences corresponding to the unique sequences or common sequences in each pool is shown in the upper graph. The points of common sequences between the pools were connected by a line. (f) The similarity between the pools was shown as a heat map. Similarity is assessed by calculating the read count ratio, which measures the ratio of common sequence read counts to the total read counts for each pool on the x-axis. (g) The read counts of the selected peptides before and after combining the variant peptides are plotted. (h) Amino acid frequencies of the naïve T-Surf library and the redundant peptides from each enriched pool were compared. The fold change of the frequency shows that the aromatic residues (Trp, Tyr, and Phe) and Leu are mainly enriched in the selected peptides.

Oligonucleotide pools of the designed library were synthesized and fused to the N-terminus of the M13 bacteriophage P8 coat protein, allowing a polyvalent peptide display on 5–40% of the total ∼2,700 copies of the P8 protein on each phage^24^. We also constructed the T-Surf library displayed on the minor coat protein P3, which resulted in a monovalent display of a peptide, to maximize the likelihood of finding TIR-binding sequences. Next-generation sequencing (NGS) of the T-Surf phage libraries confirmed that at least 99.7% and 54.8% of the designed peptides were present in the P8 and P3 libraries, respectively (**Fig. 1b** and **Supplementary Table 3**). While the designed peptides were the major population in both pools (**Fig. 1c**), 997,373 and 603,588 peptide variants including any amino acid substitution in the correct frame were also included in the final T-Surf P8 and P3 libraries during the library construction; so the arithmetic library diversity used for the screening should be much higher—1,187,792 and 708,160 for T-Surf P8 and P3, respectively (**Fig. 1d** and **Supplementary Table 3**).

### Selection of MAL– and MyD88-binding peptides

T-Surf libraries were screened against the immobilized TIR domains of two major human TLR adaptor proteins, MAL^TIR^ and MyD88^TIR^, allowing cross-species selection of TIR-binding motifs with 190,946 TIR-derived peptides in a single binding assay (**Fig. 1a**). Enriched phage pools from the fourth round of selection were analyzed using NGS after barcoding the peptide-coding regions of the phage pools. For each enriched pool, the most redundant 500 peptide sequences are summarized in **Dataset 2**, and the origin and the structural region of the top 11 peptides from the pools are described in **Dataset 3**. When the enriched peptides from the pools (P8_MAL, P8_MyD88, P3_MAL, and P3_MyD88) were compared, no significant similarity was observed (**Fig. 1e,f**) indicating the T-Surf P3 library selection is complementary to that of the P8 library, presumably owing to the different valency of the display^24^. Rather, a small fraction of overlapping sequences was found in both P3_MAL and P3_MyD88, indicating promiscuous binding of TIR-derived peptides. Although there were a few non-designed peptides with a point mutation compared to the originally designed sequences in the enriched population, these variants were scarce compared to the designed peptides, as the ranking of the selected peptides remained similar when we counted the original and the variants (containing a point mutation) as a single representative peptide (**Fig. 1g** and **Dataset 4**). Furthermore, the redundant peptides with an excess population >0.015% in the naïve T-Surf libraries were not enriched (**Supplementary Tables 4** and **5**), indicating that the abundance of the peptides did not inherently bias the selection against the targets. Notably, in our screening, the peptides from human TIRs or the reported decoy peptides^11^ did not outcompete the newly discovered sequences. This confirmed that the T-Surf library is a potent and economical tool for screening TIR-derived TIR-binding motifs in parallel, compared with previous approaches that used a few synthetic peptides fragmenting the human TIR domains or bacterial/viral TIR-binding domains^15–20^. The structural region of the redundant peptides (**Dataset 3**) varied from R2 to R11, among which R2, R9, and R11 were the major ones. The peptides originating from the same structural region showed some degree of sequence similarity, with a few exceptions to define a consensus motif (**Supplementary** Fig. 2). Moreover, aromatic residues (Trp, Tyr, and Phe) and leucine were prevalent in the enriched peptide pools compared with those in the naïve T-Surf library (**Fig. 1h**).

### Suppression of TLR signaling by synthetic peptides

To evaluate the TLR-inhibitory effect of the selected TIR-binding motifs, 21 peptides (TIR-derived immunomodulatory peptides; TDIPs) and a negative control peptide (CP) were synthesized, including the cell-penetrating peptide (CPP) penetratin, at the N-terminus for the cell membrane permeability (**Fig. 2a,b**). Effective intracellular delivery by penetratin has been confirmed in previous studies on the design of cell-permeable decoy peptides targeting TLRs^15–20^. The immunomodulatory effects of TDIPs, exerted by interfering with TLR signaling, were first evaluated on HEK-Blue^TM^ hTLRs by measuring SEAP signals. The TDIPs exhibited a broad range of TLR inhibition, especially against TLRs 2, 4, 7, and 9 (**Fig. 2c**). After confirming the negligible cytotoxicity of TDIPs except TDIP4, 8, and 10 (**Fig. 2d**), we confirmed the dose-dependent inhibitory effect of TDIPs on TLR4 signaling (**Fig. 2e**). Based on the results, the most potent four TDIPs were selected for further investigation: TDIP1, 5, 6, and 20.

**Figure 2.**
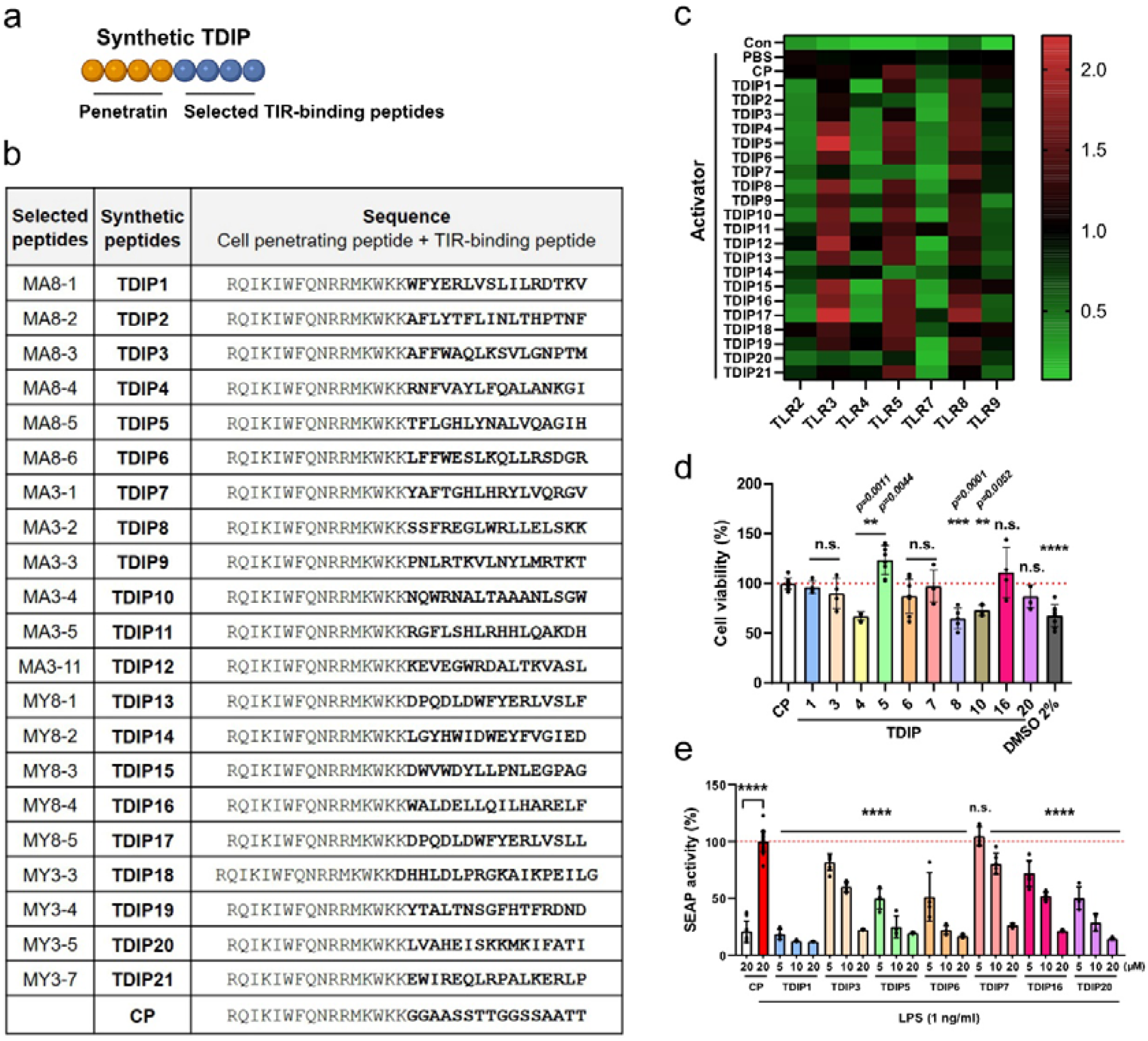
Suppression of TLR signaling by the selected peptides. (a) The selected peptides were synthesized with a cell-penetrating peptide, penetratin, at the N-terminus. Created with BioRender.com. (b) The full sequences of selected peptides were displayed. (c) The TLR inhibitory effects of the TDIPs evaluated on the HEK-Blue^TM^ hTLRs were displayed as a heat map (n = 3). Cells were activated with 100 ng/ml Pam3CSK4 (on TLR2), 10 μg/ml poly(I:C) (on TLR3), 100 ng/ml LPS (on TLR4), 10 ng/ml FLA-ST (on TLR5), 5 μg/ml imiquimod (on TLR7), 1 μg/ml R848 (on TLR8), or 10 μg/ml ODN2006 (on TLR9). (d) Cytotoxicity of the peptides was assessed using HEK-Blue^TM^ hTLR4 (n = 4–9). Data are presented as mean ± SD. The *P* values were calculated by one-way ANOVA. All significance is versus CP group. (e) D ose-dependent inhibition of TLR4 signaling by TDIPs was evaluated by measuring SE AP activity in the HEK-Blue^TM^ hTLR4 (n = 4–24). Data are presented as mean ± S D. *\*\*\*\*p < 0.0001*. All significance is versus LPS-treated CP group. n.s., not signific ant.

Next, we investigated the anti-inflammatory action of the selected TDIPs on RAW264.7, a mouse macrophage cell line. Although the peptides were selected against human TIRs, the TIR domains of human and mouse MAL and MyD88 are highly conserved as in other mammalian species, unlike the extracellular domains of TLR4 that differ considerably between species, probably owing to species-specific adaptation to microbial environments^32^. Therefore, we hypothesized that the peptides might be effective in mouse cell lines. TLR4 activates NF-κB through either the MyD88-or TRIF-dependent pathway, with phosphorylation of IRAK4 or IRF3, respectively, serving as indicator of pathway activation.

Furthermore, mitogen-activated protein kinase (MAPK) activation is a prominent downstream pathway of MyD88. First, the selected TDIPs significantly suppressed IRAK4 and IRF3 phosphorylation, indicating that both MAL-MyD88 and TRAM-TRIF signaling were abrogated by all the TDIPs (**Fig. 3a**). The selected TDIPs also inhibited ERK, JNK, and p38 phosphorylation in a dose-dependent manner (**Fig. 3b,c**). Furthermore, the regulatory effect of the peptides on NF-κB was confirmed by analyzing the phosphorylation of both IκBα and NF-κB, and NF-κB nuclear translocation; treatment with TDIPs reversed the effects of both LPS-induced phosphorylation of IκBα and NF-κB, and IκBα degradation (**Fig. 3d,e**), indicating the inhibitory effect of TDIPs on NF-κB activation. Simultaneously, the nuclear translocation of NF-κB after LPS treatment was also blocked by TDIPs (**Fig. 3f**).

**Figure 3.**
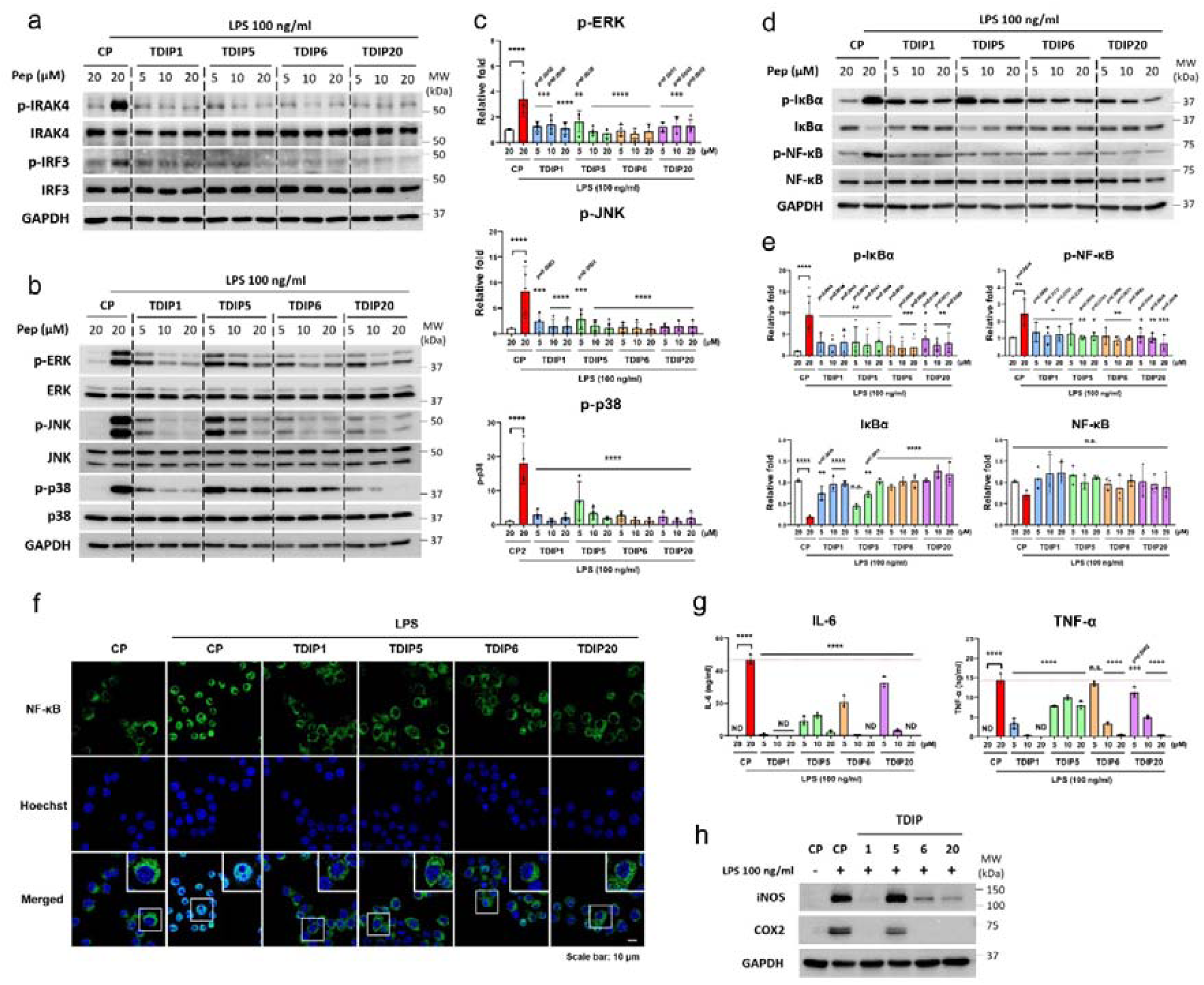
TDIPs block TLR4 signaling in mouse macrophages. (a) Immunoblot analysis of IRAK4 and IRF3 phosphorylation. (b, c) Immunoblot analysis and quantification of p-ERK (n = 4), p-JNK (n = 3–5), and p-p38 (n = 4). (d, e) Immunoblot analysis and quantification of p-IκBα (n = 4–5), IκBα, p-NF-κB, and NF-κB (n = 3). (f) Immunostaining for NF-κB (green) with Hoechst (blue) staining. Scale bar = 10 μm. (g, h) RAW264.7 cells treated with TDIPs for 1 h and followed by treatment with LPS for 24 h. ELISA (g) of IL-6 and TNF-α (n = 3), and immunoblot analysis of iNOS and COX2 (h). Data are presented as mean ± SD. *\*\*\*\*p < 0.0001*. All significance is versus LPS-treated CP group. n.s., not significant. ND, not detected.

Furthermore, the inactivation of NF-κB by TDIPs led to a decrease in proinflammatory cytokine levels; both IL-6 and TNF-α secretion was significantly attenuated by TDIPs in a dose-dependent manner (**Fig. 3g**). We further confirmed that TDIP1, 6, and 20 decreased iNOS and COX2 expression (**Fig. 3h**). Overall, these findings demonstrated that TDIPs block LPS-induced inflammatory TLR4 signaling.

### Therapeutic efficacy of TDIPs in mouse model of LPS-induced sepsis

In sepsis, stimulation of proinflammatory cytokines by bacterial components such as LPS often leads to a severe cytokine storm and death. Thus, we tested the anti-inflammatory potential of TDIPs by inhibition of TLR4 in mouse sepsis model. To evaluate the therapeutic effects of TDIPs on LPS-induced septic shock, we first examined the survival rate of TDIP-treated mice after intraperitoneal LPS challenge (30 μg/g). Mice treated with LPS and a vehicle (PBS) experienced total mortality, and mice treated with LPS and the CP exhibited a 20% survival rate (**Fig. 4a**). In contrast, mice treated with TDIPs (TDIP1, TDIP5, TDIP6, and TDIP20) showed significantly higher survival rates (100%) than mice treated with the vehicle or CP (p < 0.01 vs. vehicle, p < 0.05 vs. CP). Since proinflammatory cytokine overproduction is indicative of sepsis, we evaluated the levels of TNF-α, IFN-γ, IL-1α, IL-1ß, IL-6, IL-10, IFN-ß, IL-17A, IL-23, IL-27, MCP-1, IL-12p70, and GM-CSF in the serum at 16 h after LPS-induced septic shock. The protein levels of these 13 proinflammatory cytokines decreased significantly in all TDIP-treated mice compared with those in the vehicle-or CP-treated mice (**Fig. 4b**). Histopathological examination of kidney tissues using H&E staining revealed significantly attenuated LPS-induced damage in the mice treated with TDIPs compared with that observed in the vehicle-or CP-treated mice (**Fig. 4c**). In addition, TUNEL assay demonstrated a reduction in the number of TUNEL-positive apoptotic cells in all TDIP-treated mice compared with that in sepsis samples (from vehicle-or CP-treated mice) (**Fig. 4c**). Overall, TDIPs provide protective effects against sepsis in mice by significantly improving the survival rate through reducing systemic inflammatory response and attenuating renal failure.

**Figure 4.**
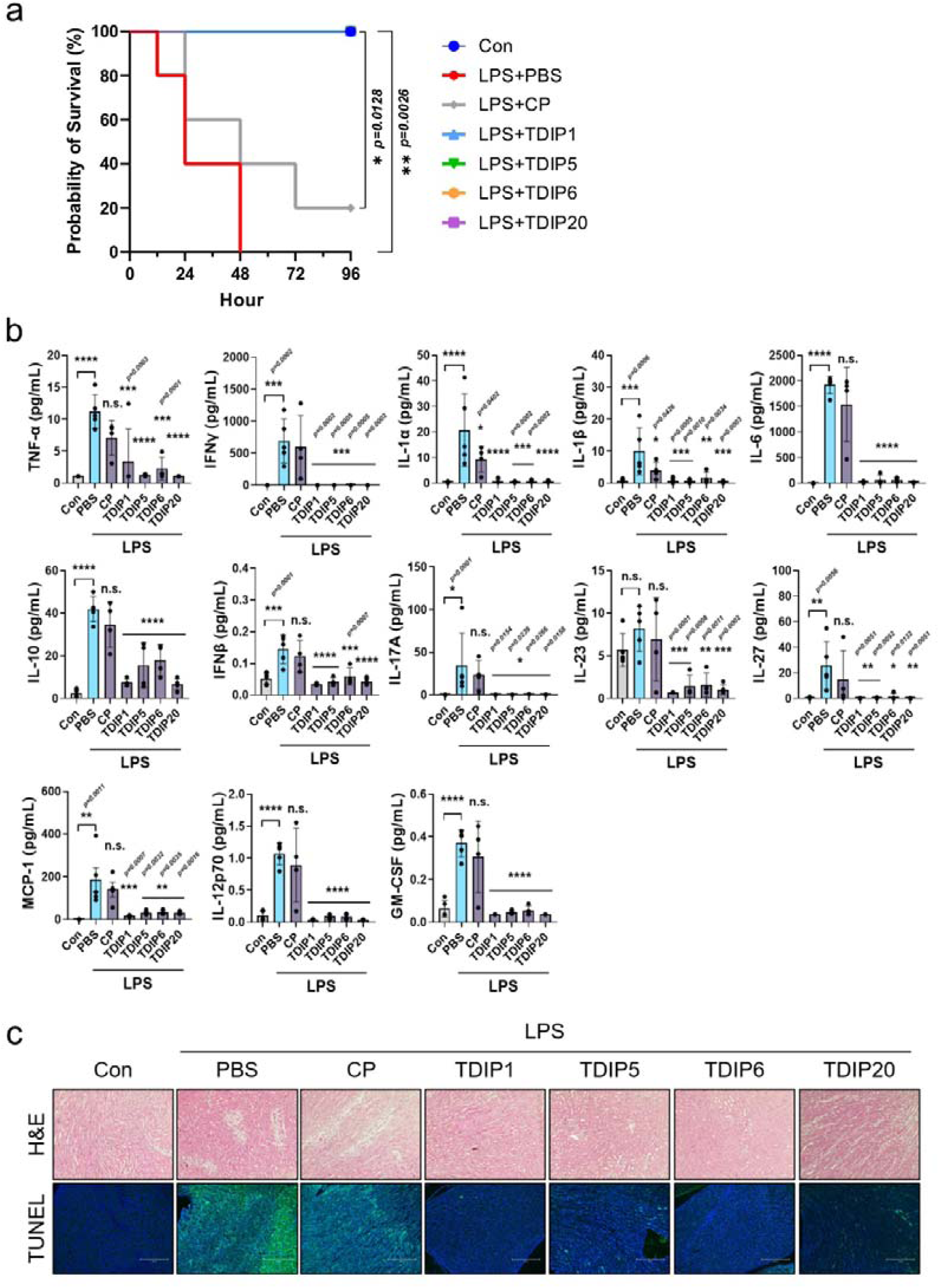
Therapeutic effect of TDIPs in LPS-induced sepsis model. (a) Survival of mice (n = 5) for 96 h after injection of LPS (30 μg/g) and treatment with CP, TDIP1, 5, 6, or 20 (10 nmol/g). Mice treated with TDIPs showed 100% survival rate as the control group. (b) Measurement of TNF-a, IFNg, IL-1a, IL-1b, IL-6, IL-10, IFNb, IL-17A, IL-23, IL-27, MCP-1, IL-12p70, and GM-CSF in serum at 16 h after the injection of LPS and indicated peptides (n = 4–5). Data are presented as mean ± SD. *\*\*\*\*p < 0.0001*. All significance is versus LPS-treated vehicle group. n.s., not significant. (c) Representative H&E-stained images (top) and TUNEL-stained images (bottom) in the kidney tissues. Scale bar = 100 μm.

### Protective effects of TDIP eye drops against sodium iodate (NaIO_3_)-induced retinal degeneration

TLRs in RPE cells are implicated in the regulation of inflammatory responses and oxidative stress, thereby influencing the progression of age-related macular degeneration (AMD). Therefore, we investigated the protective effect of TDIPs against NaIO_3_-induced retinal degeneration in a mouse model paralleling the human condition of dry AMD characterized by retinal dysfunction, drusen-like deposits, and atrophy of RPE cells and photoreceptors^33^. Upon being administered NaIO_3_ along with a vehicle or CP, the mice exhibited numerous yellow-white, drusen-like structures in their fundus (**Fig. 5a**). Optical coherence tomography (OCT) further revealed significant alterations in the RPE layer, including an abundance of high reflex zones and noticeable thinning of the retinal structure. Contrastingly, mice treated with NaIO_3_ alongside TDIP1 or TDIP5 eye drops showed a significant decrease in these pathological features. Histological analysis using H&E staining revealed extensive melanin accumulation in the RPE layer of mice treated with NaIO_3_ and a vehicle or CP, but not in that of the normal group (Con). The structural integrity of the photoreceptor inner segment and outer segment was damaged in both the vehicle– and CP-treated groups, resulting in retinal folding and disruption (**Fig. 5a**). However, TDIP1 or TDIP5 treatment significantly improved retinal morphology, as seen by the maintenance of total retinal and ONL thickness close to normal levels (**Fig. 5b,c**). TUNEL staining revealed elevated photoreceptor cell death in the vehicle– and CP-treated groups but a significant decrease in cell death in all TDIP-treated mice relative to the vehicle group (**Fig. 5d**).

**Figure 5.**
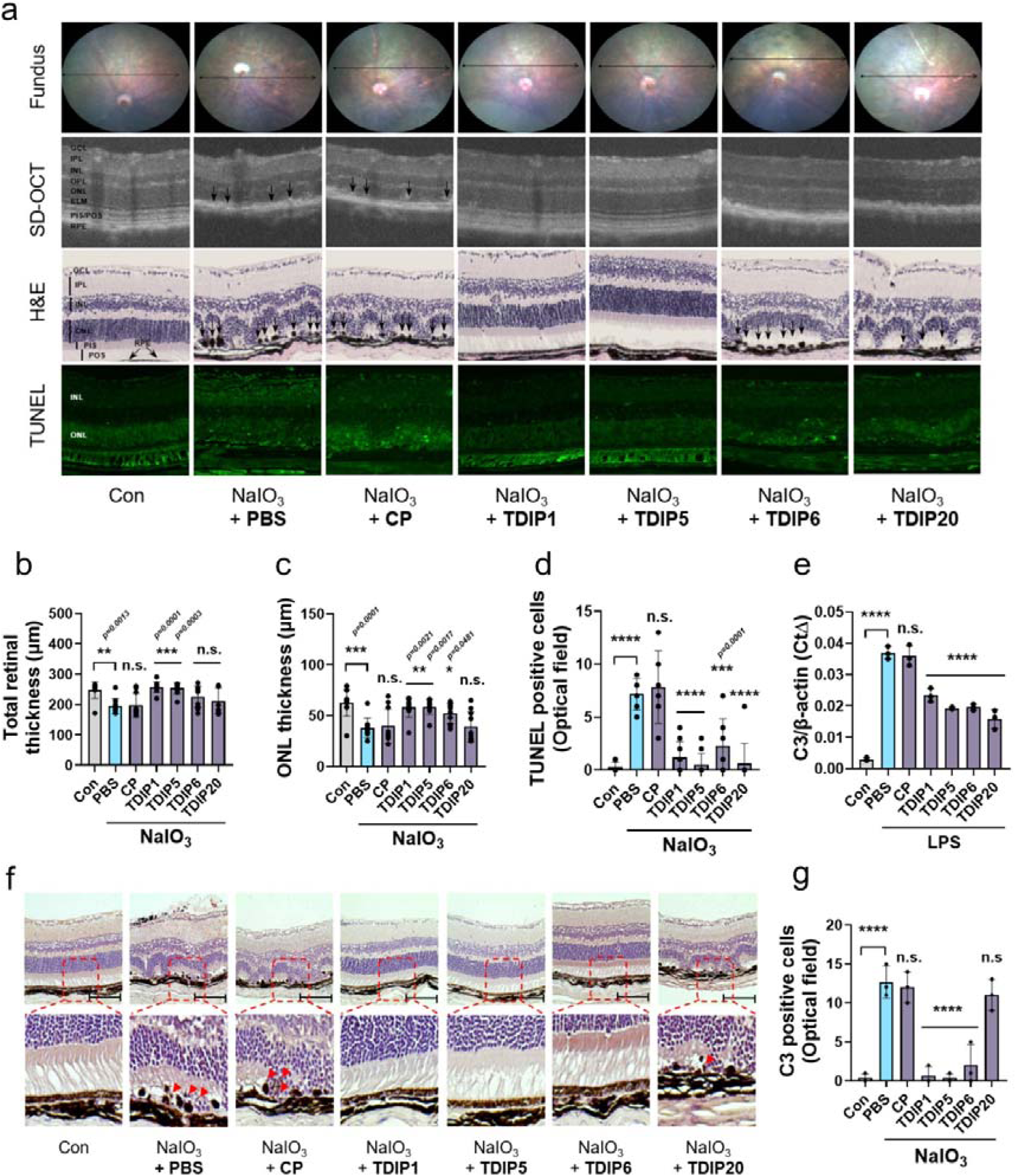
Therapeutic effect of TDIPs on NaIO_3_-induced histological changes and C3 expression in mouse retina. (a) Representative images of fundus, SD-OCT, H&E, and TUNEL staining showing the therapeutic effect of TDIPs. (b-d) Total retinal thickness, ONL layer thickness and TUNEL-positive cells of NaIO_3_– and TDIPs-treated mice are compared (n = 6–10). (e) C3 mRNA in BMDM cells was measured by quantitative real-time RT-PCR. (f) Immunohistochemical staining shows the decrease of C3 in TDIPs-treated retina. (g) C3-positive cells in retina were counted. Data are presented as mean ± SD. *\*\*\*\*p < 0.0001*. All significance is versus NaIO_3_-induced vehicle group. n.s., not significant.

We further investigated the mechanism underlying the protective effects of TDIPs by focusing on the modulation of complement factor 3 (C3) expression, a key element in TLR signaling and photoreceptor cell death^10^. We first treated LPS-primed bone marrow-derived macrophages (BMDMs) with TDIPs to evaluate the effect of the peptides on C3 expression. LPS-exposed BMDMs showed a significant increase in C3 expression. However, treatment with all TDIPs reduced C3 expression (**Fig. 5e**). Immunohistochemical staining for C3 revealed a higher frequency of C3-positive cells in the vehicle and CP groups compared to the normal retina, but eye drops of TDIP1, 5, and 6 markedly decreased C3-positive cells compared with that in the vehicle-treated group (**Fig. 5f,g**). Overall, TDIPs effectively suppressed the TLR-mediated upregulation of C3, thereby protecting the RPE and reducing photoreceptor cell death under NaIO_3_-induced retinal degeneration conditions.

### Molecular characteristics of selected TDIPs

To better understand the molecular properties of TDIPs, a dot blot immunoprecipitation assay was performed to identify the potential binding of TDIPs to the other TIR domains of adaptor proteins. FLAG-tagged TIR domains of MAL, MyD88, TRAM, and TRIF were expressed in HEK293T cells and immunoprecipitated from cell lysates supplemented with TDIPs. Four TDIPs and one CP were tested against each TIR domain. Co-immunoprecipitation confirmed a robust interaction between the TDIPs and four TIR domains (**Fig. 6a**). In contrast, CP did not bind with MAL^TIR^, MyD88^TIR^, and TRAM^TIR^. TRIF^TIR^ showed a non-specific interaction with all peptides. This suggests that the multifaceted inhibitory effect of TDIPs on multiple TLRs is a result of their multispecific binding to TIR domains.

**Figure 6.**
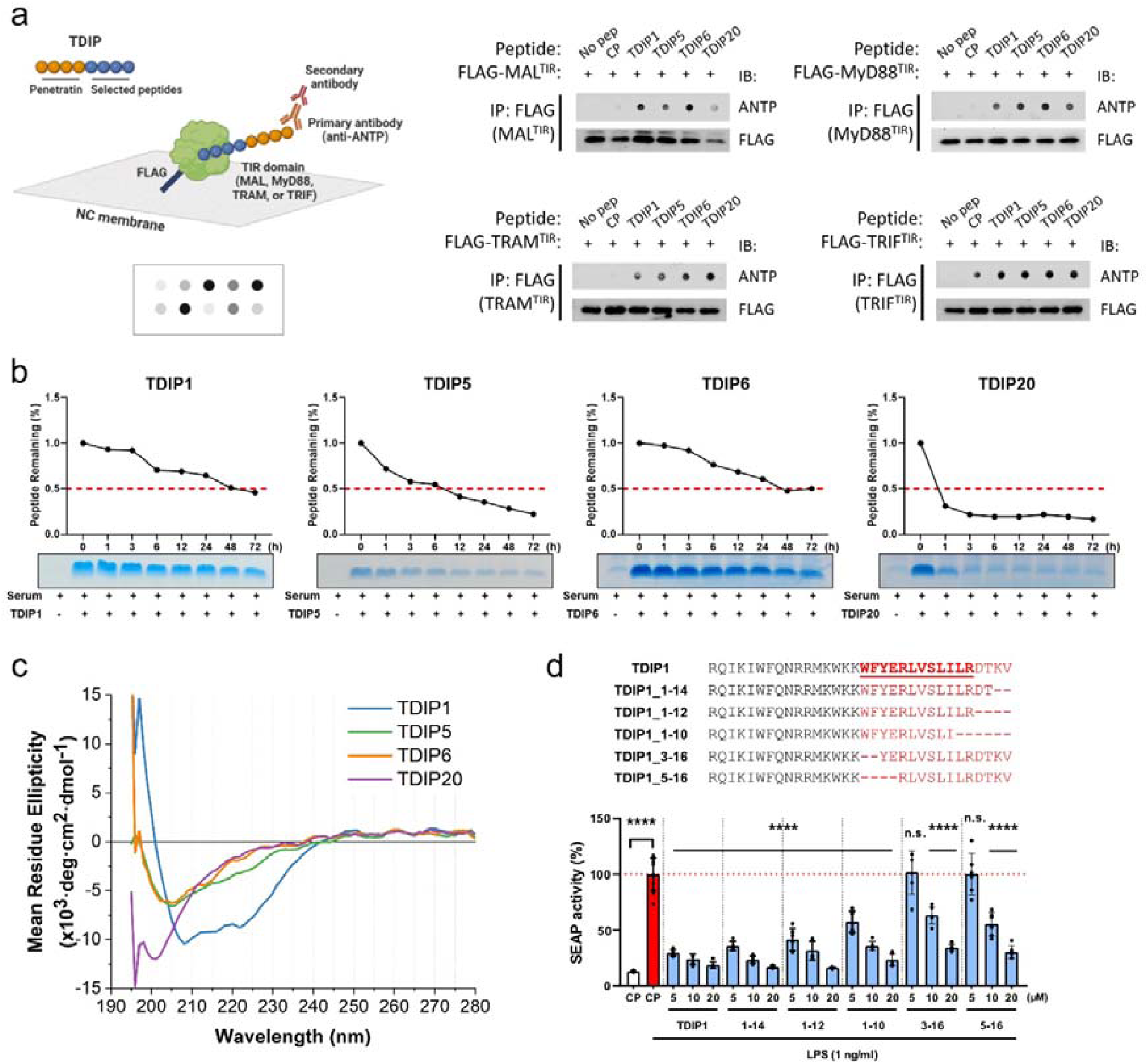
Molecular characterization of TDIPs. (a) Co-immunoprecipitation of TIR domains and TDIPs. Lysates of HEK293T cells expressing FLAG-tagged TIR domains were incubated with the indicated TDIPs. Peptides were blotted with an antibody against the penetratin sequence. Created with BioRender.com. (b) Human serum stability (half-life) of the TDIPs was measured. The percentage of remaining peptide was quantified up to 72 h. (c) The secondary structure of the selected TDIPs was determined by circular dichroism. While TDIP1 retains the helicity (minimum at 208 nm and 222 nm), other TIDPs show a more disordered structure. (d) TDIP1 and its truncates were compared for inhibition of TLR4 signaling by using HEK-Blue^TM^ hTLR4 cells. Data are presented as mean ± SD. *\*\*\*\*p < 0.0001*. All significance is versus LPS-treated CP group. n.s., not significant.

We next measured the human serum stability of the four TDIPs. After TDIPs were exposed to serum proteases by incubation with human serum, the remaining intact peptides were quantified in a time-dependent manner. Four peptides showed different half-lives in human serum (**Fig. 6b**). In particular, TDIP1 and TDIP6 exhibited high stability, considering that linear peptides often have a half-life of several minutes to a few hours; up to 50% of the remaining peptides were detected even after 48 h of incubation. While TDIP5 had a half-life of around 6 h, TDIP20 had a shorter half-life of <1 h. We also measured circular dichroism (CD) spectra of the four TDIPs to evaluate the secondary structure. While TDIP1 retains the helicity (minimum at 208 nm and 222 nm), the spectra of the other TDIPs are typical of unstructured peptides (**Fig. 6c**).

To identify the minimal functional residues of TDIP1, we then compared the TLR4-inhibitory activity of five truncated forms of TDIP1 in LPS-primed HEK-Blue^TM^ hTLR4 cells by measuring SEAP signals. As a result, the C-terminal four residues of the 16 amino acid-binding moiety of TDIP1 were identified as not critical for inhibition of TLR4 signaling, whereas the N-terminal hydrophobic amino acids are the key residues (**Fig. 6d**), which will guide further optimization of the efficacy of TDIPs.

## Discussion

We established a comprehensive screening platform for identifying TIR-binding motifs that can be exploited for immunomodulation in inflammatory diseases. This peptide fragment-based high-throughput screening, recently enabled by advances in parallel oligo-synthesis and NGS analysis, has been widely utilized to map functional regions of parent proteins interacting with a partner protein^24,29,31^. In this study, the proteomic TIR-derived custom peptide library successfully uncovered TIR-binding motifs from homologous TIR domains of unprecedented species. We found that the source domains of the selected peptides were highly diverse in origin (**Dataset 3**) and that the sequences found were distinct from previously reported TLR-inhibitory peptides from the human TIR domain^11^. Notably, we discovered R2-derived peptides as major TIR-binding fragments in addition to R9 and R11 from our screening platform. While the surface regions from R3 to R11 of several TIR domains are major participants in TIR-TIR interactions, R2 has rarely been considered as a potential inhibitory peptide^11^. The enrichment of four amino acids – Trp, Tyr, Phe, and Leu – in the selected peptide pools from the screening is consistent with the previous observation on the types of amino acids appearing as hotspot residues in the helical interfaces^34^, considering that TIR-TIR interactions are mainly formed by α-helices surrounding the TIR surface. Taken together, we have confirmed that our screening method is a highly effective way to explore TIR-binding sequences from the vast untapped TIR domains.

With the data from the selections, we demonstrated that the TIR-binding peptides effectively regulate TLR-mediated signaling, exhibiting therapeutic benefits in *in vivo* models. Extended screening of the T-Surf library against other human TIR-containing proteins (TLRs, TRIF, and TRAM) and homologous TIR domains from pathogens could help identify suitable peptides applicable to, for instance, infectious diseases and immune disorders. Additionally, cell-based phenotypic screening will unlock the full potential of the T-Surf library in modulating TLR pathways. The newly discovered TIR-binding peptides against other TIRs, in addition to the TDIPs described in this study, will provide an extensive resource for TLR modulation.

We demonstrated the efficacy of the selected TDIPs against LPS challenge both *in vitro* and *in vivo*. Treatment of macrophage cells with synthetic TDIPs containing a cell-penetrating moiety suppressed TLR4-dependent inflammatory pathways in a dose-dependent manner. Moreover, the selected TDIPs exerted broad-spectrum TLR-inhibitory effects by inhibiting interactions between TIR domain-containing proteins, which enhanced the therapeutic efficacy in the murine sepsis model. Additionally, we demonstrated the protective effects of the selected TDIPs in a preclinical model of dry AMD. Although two drugs for dry AMD (pegcetacoplan and avacincaptad pegol) have recently been approved by the FDA, TDIPs differ in their mechanism of action and can be administered as eye drops. In the exploration of therapeutic strategies for AMD, inhibition of TLR signaling has emerged as an important mechanism. This is because TLR signaling plays a critical role in triggering the expression of C3, which is a major contributor to the development of AMD^10^ and a target protein of pegcetacoplan. The results of our study show that TDIPs can inhibit the progression of AMD by limiting TLR signaling and thereby inhibiting the expression of C3. The topical application of TDIPs can ensure dosing convenience for patients. Intravitreal injection of therapeutic agents is associated with serious side effects and various ocular complications^35^; the noninvasiveness of TDIP eye drops is expected to mitigate such complications. Although additional studies are needed to elucidate the delivery and efficacy of TDIPs in large animal models and humans, as previously described^36^, penetratin exhibited excellent cellular uptake in human conjunctival epithelial cells and rabbit cornea, with lower cellular and tissue toxicity than that of other CPPs such as TAT (Transactivating transcriptional activator) and polyarginine. The penetration efficiency of penetratin can be further improved by optimizing the sequence for better efficacy, without increasing toxicity^37^. Thus, penetratin-mediated delivery of TDIPs via topical administration may hold promise as a novel treatment modality for dry AMD.

Many essential cellular processes depend on the protein interactome, and deregulation of the PPI network is often associated with the pathology of various diseases. However, the vast intracellular PPI network remains an untapped area for drug development, as it is not an optimal target for regulation by major drug modalities involving small molecule-or antibody-based targeted therapies. To this end, peptides with suitable physicochemical properties against intracellular PPIs are receiving increasing attention^38–40^. Protein fragment-based high-throughput screening has been widely adopted to reveal minimal sequence regions sufficient to intervene in an interaction with a binding partner, through which a handful of peptides with therapeutic potential have been discovered^30,31^. As such, our strategy can be readily adapted by diverse studies in which multiple homologous domains mediate interactions over short surface segments. We found that functional domains participating in signal transduction serve as a promising source of bioactive peptides that modulate intracellular PPIs.

The general application of the T-Surf library selection has certain limitations. The unavailability of target proteins, mainly because of insoluble expression in *E.coli* such as TRIF^TIR^ and TLR4^TIR^ in our attempt, hampered the phage display selection against the target TIR domains. This may be resolved by employing cell-free protein synthesis^41^ or mammalian expression. Regarding the therapeutic application of TDIPs, the efficacy and safety of TDIPs could be optimized at the amino acid level, based on the structural information of the TIR-peptide complex. Since the helicity of an α-helical peptide largely affects on its activity and stability^42^, the helicity of TDIPs can also be optimized to improve the potency and stability of the peptides using several approaches, including the cyclization of peptides using non-canonical amino acids or chemical crosslinkers^43^. Indeed, although the selected peptides were predicted by PEP-FOLD4^44^ to have helical conformations similar to that of their original TIR domains (**Supplementary** Fig. 3), CD spectra of the three TDIPs out of four showed low level of helicity in contrast to the original TIR domain or the predicted α-helical conformation.

This is not surprising since α-helical peptides in a short linear form lack factors that stabilize their folding^43,45^. Additionally, various strategies such as enhancing cellular internalization and stability, or reducing immunogenicity, can be applied to maximize the efficacy of TDIPs and overcome potential challenges in TDIP-based treatments^46^. The off-target binding of TDIPs must be investigated to control any unexpected side effects in *in vivo* applications. Further in-depth studies are warranted to elucidate whether the multifaceted effect of multispecific binding of TDIPs is critical for the *in vitro* and *in vivo* efficacy of TDIPs.

## Materials and methods

### Design of TIR-derived peptide library

The protein sequences homologous to the TIR domain were retrieved from a full sequence alignment for the TIR domain (PF01582) in the Pfam database, which consists of 13,644 sequences (as of March 2021). The solution structure of the TIR domain of human MyD88 (PDB ID: 2Z5V) was manually inspected, and nine (2∼11) surface-exposed regions were identified (**Supplementary** Fig. 1). The alignment blocks corresponding to these selected regions (Residue index 21-36, 37-52, 48-63, 56-71, 77-92, 89-104, 110-125, 119-134, and 133-148 in 2Z5V) were consolidated and then fragmented into non-redundant 16 amino acid peptides without a gap (with a sliding window of size 1). In cases where peptide fragments were shorter than 16 amino acids, we extended them by including adjacent residues to ensure uniformity in length. Consequently, we generated a collection of 779,406 unique peptide sequences derived from the TIR domain. To streamline the TIR-derived peptide library, we performed sequence clustering with 82% sequence identity. Moreover, the sequences of 15 human TIR domains and 4 putative TIR-binding domains (**Supplementary Table 1**) were included in the final list, resulting in a refined library containing 190,945 sequences. Next, every cysteine was substituted with alanine to avoid potential complications arising from free cysteines in the phage coat proteins. Subsequently, these peptide sequences were back-translated with optimizing codons by matching the codon usage of *E. coli*. Two DNA fragments (AATGCCTATGCAGCATCTTCATCTGGATCCATGGCT and CTCGAGGGTGGAGGATCTGGAGGAGGTGCAGAAGGT) were added to the 5’ and 3’ ends, respectively, to anneal the primers and subsequently clone them into the P8 and P3 phagemids. The final oligonucleotide pool was obtained from Twist Bioscience (San Francisco, USA).

### Construction of T-Surf phage library

The oligonucleotide library was amplified with a pair of primers using PrimeSTAR® Max DNA Polymerase (Takara Bio). The P8 and P3 phagemids were double-digested using NcoI-HF® (NEB) and XhoI (NEB). The library amplicons (7 ng) were each mixed with 210 ng of linearized P8 and P3 phagemids, and cloning reactions were performed at 50 °C for 1 h using the In-Fusion® HD Cloning Kit (Takara Bio). The reaction mixtures were further transformed into TG1 electrocompetent cells (Lucigen). The cells were rescued with 1 ml of recovery medium for 1 h, followed by the addition of 1 μl M13KO7 helper phage (NEB).

After incubation for 30 min, the cells were transferred to 30 ml of 2×YT medium (16 g Bacto tryptone, 10 g Bacto yeast extract, and 5 g NaCl in 1 l water) containing carbenicillin (100 μg/ml) and kanamycin (25 μg/ml) and grown for 16 h at 37 °C, with shaking at 200 rpm. The cultivated cells were concentrated via centrifugation at 12,000 ×g for 10 min, and the phage particles in the supernatant were precipitated by adding 1/5 volume polyethylene glycol (PEG)–NaCl (20% PEG-8000 (w/v) and 2.5 M NaCl), followed by incubation for 10 min at 4 °C and centrifugation at 16,000 ×g at 4 °C for 20 min. The phage pellet was resuspended in 2 ml phosphate-buffered solution (PBS) with 0.05% Tween-20 and 0.5% bovine serum albumin (BSA) and further centrifuged at 5,000 ×g for 2 min to remove insoluble debris. The final concentration of the phage library was determined by measuring the absorbance at 268 nm using a NanoPhotometer N60 (IMPLEN).

### Protein expression and purification

Twist Bioscience synthesized the 6×His-tagged TIR domains of MAL (residues 79-221) and MyD88 (residues 148-296) as clonal genes in pET-29b(+). The plasmids expressing TIR domains were transformed into *E. coli* BL21 (DE3) cells and cultivated in LB (Luria-Bertani) medium (5 g yeast extract, 10 g tryptone, and 10 g NaCl in 1 l water). Protein expression was induced by treatment with 0.5 mM isopropyl β-d-1-thiogalactopyranoside at 37 °C for 4 h.

The cells were precipitated via centrifugation at 10,000 ×g for 5 min, and lysed through sonication in Tris-HCl (pH 8.0) containing 150 mM NaCl and 10 mM imidazole. The soluble proteins were purified using His60 Ni Superflow Resin (Takara). The purified proteins were further isolated using Superdex 75 16/600 (Cytiva), a size exclusion column.

### T-Surf library selections

Phage display screening was performed by coating two wells of a 96-well plate (Maxisorp, Thermo Fisher Scientific) with 100 μl of 5 μg/ml TIR domains overnight; then the coated wells were blocked with 200 μl of blocking buffer (0.2% BSA in PBS) and set aside for 1 h at 4 °C with gentle shaking. Next, 100 μl of the phage library was added to the wells and incubated for 1 h at room temperature (RT; 20 – 22°C) with gentle shaking. The plate was washed four times with PBS containing 0.5% Tween-20; the bound phages were eluted by adding 100 μl of 100 mM HCl to each well. The elute was neutralized with 10 μl of 1.0 M Tris-HCl (pH 11.0). *E. coli* TG1 cells were grown to the log phase (A600 = ∼0.6), and 1 ml of the culture was incubated with 100 μl of the eluted phages for 30 min at 37 °C, with shaking at 200 rpm. Next, 1 μl of M13KO7 helper phages was added, followed by incubation for 45 min at 37 °C, with shaking. Subsequently, the cells were transferred to 25 ml of 2× yeast extract tryptone/carbenicillin/kanamycin medium and incubated overnight at 37 °C with shaking at 200 rpm. The amplified phages were precipitated with 20% PEG-8000/2.5 M NaCl to be used for the next round of selection.

### NGS analysis of the peptide library

The naïve T-Surf libraries in P8 and P3 and the enriched phage pools after four rounds were barcoded for NGS as previously described^47^. Briefly, 5 μl of the phage pools was used as a template for 20 cycles of a 50 μl PCR reaction using 0.5 mM each of forward and reverse barcoded primers and PrimeSTAR® Max DNA Polymerase. The PCR products were purified using Expin Gel SV (GeneAll). The NGS samples were quantified using qPCR, following the qPCR Quantification Protocol Guide (KAPA Library Quantification kits for Illumina Sequencing Platform), and qualified using the TapeStation High Sensitivity D1000 ScreenTape (Agilent Technologies). The indexed libraries were further sequenced by Macrogen (Seoul, Republic of Korea) using the NovaSeq platform (Illumina) with 100 bp paired-end reads. The reads were demultiplexed by barcodes and filtered for an average Phred quality score of at least 30, followed by trimming the flanking consensus sequences using Cutadapt (version 2.3). Peptide sequence populations were analyzed according to the read counts.

### Peptide synthesis and reconstitution

The peptides were synthesized by ANYGEN (Gwangju, Republic of Korea) at >95% purity. The lyophilized peptides were reconstituted in ultrapure water (Invitrogen) to a concentration of 2 mM. Next, the peptide solutions were diluted with 10× PBS to make working concentrations, and added to cell culture media at concentrations mentioned for the respective *in vitro* experiments.

### Cell viability assay

HEK-Blue™ hTLR4 (InvivoGen) cells were seeded at a density of 1 × 10^5^ cells per well in a 96-well plate (Costar). After 24 h of incubation to allow adherence and growth, the cells were treated with 20 μM of TDIPs and incubated for an additional 24 h. EZ-Cytox reagent (ITSBio) was added to each well, and the plate was incubated at 37 °C for 30 min. The absorbance was measured at 450 nm using using the Multiskan SkyHigh Spectrophotometer (ThermoFisher Scientific).

### Secreted embryonic alkaline phosphatase (SEAP) activity assay

HEK-Blue™ hTLR2/1, hTLR3, hTLR4, hTLR5, hTLR7, hTLR8, and hTLR9 (InvivoGen) cells were seeded in a 96-well plate (Costar) at a density of 1 × 10^5^ cells per well. After 24 h of incubation, the cells were pretreated with various concentrations of TDIPs for 1 h, followed by treatment with 100 ng/ml Pam3CSK4 (on TLR2/1), 10 μg/ml poly(I:C) (on TLR3), 100 ng/ml LPS (on TLR4), 10 ng/ml FLA-ST (on TLR5), 5 μg/ml Imiquimod (on TLR7), 1 μg/ml R848 (on TLR8), or 10 μg/ml ODN2006 (on TLR9) for 24 h. The cell culture supernatants (20 μl) were transferred to a new 96-well plate (Costar, 3370) and mixed with 180 μl of pre-warmed Quanti-Blue Solution (InvivoGen). After 1 h of incubation in the dark, the absorbance was measured at 620 nm using the Multiskan SkyHigh Spectrophotometer (ThermoFisher Scientific).

### Western blotting and protein quantification

The cells were lysed in RIPA buffer (ThermoFisher Scientific) containing halt protease & phosphatase inhibitor cocktail (ThermoFisher Scientific). After 10 min of incubation on ice, the lysates were centrifuged at 16,400 ×g for 15 min at 4 °C and the protein concentration in the supernatants was measured using a Pierce BCA protein assay kit (ThermoFisher Scientific). Total protein (10 µg) was loaded on a sodium dodecyl sulfate (SDS)-polyacrylamide gel and blotted onto polyvinylidene fluoride (PVDF) membranes (Millipore), and set aside for 60 min at 100 V/cm, in a transfer buffer containing 25 mM Tris, 192 mM glycine, and 20% ethanol. The membrane was blocked using 5% skim milk for 1 h. After blocking, the membrane was incubated with specific primary antibodies overnight, at 4 °C.

After three washes with 1× tris-buffered saline with Tween 20 (TBST) buffer (T&I), the membrane was incubated with secondary antibodies and visualized using enhanced chemiluminescence (ECL) solutions (Promega, W1001 and Thermo Fisher Scientific, 34095), according to the recommended procedure. Protein detection was performed using iBright CL1000 (Invitrogen). Obtained images were analyzed using the ImageJ 1.54 (National Institutes of Health), and the level of each protein was quantified and normalized to levels of GAPDH or α-tubulin.

### Enzyme-linked immunosorbent assay (ELISA)

RAW264.7 cells were seeded in a 96-well plate (Costar) at a density of 5 × 10^4^ cells per well. After 24 h of incubation, the cells were pretreated with various concentrations of TDIPs for 1 h, followed by treatment with LPS for 24 h. The cell culture supernatants were collected and centrifuged at 500 ×g for 5 min at 4 °C. The supernatants were transferred to a new eppendorf tube and used for measuring the levels of cytokines using the Mouse TNF-alpha Quantikine ELISA kit and the Mouse IL-6 Quantikine ELISA kit (R&D systems), following the manufacturer’s instructions.

### Confocal microscopy

RAW264.7 cells were grown on coverslips coated with poly-L-lysine (Sigma), fixed in 10% formalin for 15 min, and permeabilized with 0.2% Triton X-100 (Sigma) in PBS for 15 min. After washing with PBS, the coverslips were incubated with primary antibody diluted with PBS for overnight at 4 °C, and further washed with PBS. Next, the coverslips were incubated with a secondary antibody for 2 h at RT. After being washed with PBS, the coverslips were mounted on microscope slides with an antifade mounting medium with 4’,6-diamidino-2-phenylindole (VECTASHIELD). Fluorescence images were examined under a confocal microscope (LSM 900, Carl Zeiss).

### Serum stability analysis

Human serum (Sigma-Aldrich, H4522) was thawed on ice and centrifuged at 18,000 ×g for 10 min at 4 °C to remove lipids. The supernatant was incubated at 37 °C for 15 min to activate the proteins. Each peptide was prepared at a concentration of 1 mM in ultrapure water and mixed with human serum in a 1:1 (v/v) ratio. The mixed solutions were incubated at 37 °C, for various periods (0, 1, 3, 6, 12, 24, 48, and 72 h). A mixture of ultrapure water and human serum was used as the blank. A total of 3 μg of the peptide was loaded on an SDS–polyacrylamide gel and stained with InstantBlue Coomassie Protein Stain (Abcam). Intensity of the remaining peptide bands were quantified using the ImageJ and plotted.

### CD analysis

CD spectroscopy was performed using a Chirascan V100 spectrometer. Peptides resolved in PBS were loaded into a 0.5 mm quartz cuvette (Hellma analytics), and the CD spectra were obtained at 195–280 nm wavelength, with a 1 nm increment. The buffer baseline was acquired using PBS and subtracted from the sample spectrum.

### Dot blotting

HEK293T cells were seeded at a density of 1 × 10^6^. After incubating for 24 h, pcDNA vectors (1 μg) containing FLAG-TIR constructs MYD88, MAL, TRAM (residues 75-235), and TRIF (residues 387-545) were transfected into the cells. After 48 h, the cells were lysed using Cell Lysis Buffer II (ThermoFisher Scientific). A total of 100 μg of protein was diluted with 500 μl of PBS and incubated with 10 μl of 1 mM peptides for 2 h at 4 °C. The 20 μl of FLAG M2 affinity gel (Sigma) was added to the mixture and incubated overnight at 4 °C. After centrifuging 15,000 × g for 3 min at 4 °C, the supernatant was discarded using a vacuum. The pellet was washed with 200 μl of Cell Lysis Buffer II and PMSF and centrifuged at 800 × g for 3 min for three times. After washing, the pellet was suspended in 15 μl of Cell Lysis Buffer II with PMSF and 35 μl of 2× sample buffer. The mixture was boiled and centrifuged at 15,000 × g for 10 min at 4 °C and the supernatant was transferred into new eppendorf tube. A total of 1 µl of supernatant was loaded onto the nitrocellulose membrane (Bio-Rad). After incubation at RT for 30 min to dry, the membrane was blocked with 5% skim milk in 1× TBST buffer for 1 h. After blocking, the membrane was incubated with the primary antibody at RT for 1 h. After three times of washes with 1× TBST buffer, the membrane was incubated with the secondary antibody for 1 h at RT. After three times of washes with 1× TBST buffer, the membrane was visualized by using ECL solution (ThermoFisher Scientific) according to the recommended procedure. ImageQuant LAS 4000 (GE Healthcare) was used for the detection of protein.

### LPS-induced sepsis model

Animal studies were performed within a pathogen-free barrier zone at Korea Institute of Science and Technology (KIST) Gangneung Institute and were conducted according to the guidelines approved by the Animal Care and Use Committee of KIST (approval number: KIST-2021-09-104). Six-week-old female C57BL/6 mice purchased from Orient Bio (Orient Bio Inc., Seongnam, Republic of Korea), weighing 20–25 g, were used in this experiment.

The mice received LPS in sterile PBS through intraperitoneal injections (n = 5 per group). In the treatment group, CP or TDIPs were administered intraperitoneally at a dose of 10 nmol/g body weight in combination with LPS (50 μg/g body weight for survival rate assessment under the influence of LPS, and 30 μg/g body weight for serum cytokine and renal apoptotic cell analysis). The control group received an equivalent volume of vehicle (PBS) along with the LPS injections. Survival status was monitored at fixed intervals. Serum samples were collected 16 h after LPS injection and stored at –80 °C until analysis. Proinflammatory cytokine concentrations were analyzed using the BioLegend LEGENDplex Mouse Inflammation Panel (13 plex) with V bottom plate, according to the manufacturer’s instructions. Data analysis was performed using the LEGENDplex Data Analysis Software. For histological analysis, kidney tissue sections were fixed in a 10% formalin solution overnight, embedded in paraffin wax, and sectioned into 4 μm slices using a microtome.

Slides were stained with hematoxylin and eosin (H&E), according to standard guidelines. Apoptotic cells were quantified via the terminal deoxynucleotidyl transferase dUTP nick end labeling (TUNEL) assay using the In situ Apoptosis Detection kit (Takara Bio). Visualization of apoptotic cells was performed using the Evos FL Cell Imaging System (Life Technologies).

### Dry AMD model

Animal experiments were conducted in accordance with the guidelines of the Association for Research in Vision and Ophthalmology Statement for the Use of Animals in Ophthalmic and Vision Research. Six-week-old female C57BL/6J mice were used for this experiment. The mice were maintained under a 12/12 h light/dark cycle before the start of the experiments.

The mice received a single intraperitoneal injection of NaIO_3_ (30 μg/g)^33^ and PBS solution. The control mice received an intraperitoneal injection of PBS only. After NaIO_3_ injection, a 5 μl of 20 μM CP or TDIPs were daily applied directly to the superior corneal surface of each eye using a micropipette for 21 d (n = 10 per group). Twenty days after NaIO_3_ administration, spectral domain optical coherence tomography (OCT) and live bright-field fundus imaging were performed using the image-guided OCT system (Micron IV, Phoenix Research Laboratories) and Micron Reveal software. Before imaging, the mice pupils were dilated with 1% tropicamide and 2.5% phenylephrine, and anesthesia was induced using Zoletil (Zoletil 50, Virbac Sante Animale, France). The eyes of the anesthetized mice were lubricated, positioned in front of the OCT camera, and imaged. A total of 20 frames were averaged along the horizontal axis over the optic disc of each eye. The thickness from the outer nuclear layer (ONL) to the retinal pigment epithelium (RPE) was measured at 1080 points along the retinal section using the InSight software. The mice were euthanized 21 d after NaIO_3_ injection. The eyes were enucleated and fixed in a 10% formalin solution overnight, embedded in paraffin wax, and sectioned into 4 μm slices using a microtome. Ocular tissue slides were stained with hematoxylin and eosin, according to standard guidelines. The slides were treated with hematoxylin buffer at RT, rinsed three times with distilled water, and immersed in a 1% eosin Y solution. Photoreceptor degeneration was quantified by counting the number of nuclei per row in the ONL. This count was performed in an area of 450 pixels in height in the center of the ONL in each image, and the area of the ONL was measured using ImageJ.

## Statistical analysis

Data from at least three independent experiments are expressed as the mean ± standard deviation (SD). Statistical significance was determined using a one-way analysis of variance followed by Dunnett’s multiple comparisons test using the GraphPad Prism 10.2.1 software (GraphPad).

## Data availability

The data supporting the findings of this study are available from the corresponding author on reasonable request.

## Acknowledgments

This work was supported by an intramural grant from the Korea Institute of Science and Technology and National Research Foundation of Korea (NRF) grant funded by the Korea government (MSIT) (2021R1C1C1003843).

## Author contributions

M.-H.S., W.-B.L. and K.P. conceived, designed and supervised the study. M.-H.S., W.-B.L., K.P., and Y.L. wrote the manuscript. Y.L., T.K.K., M.I.K., and J.-Y.K. performed most of the experiments. D.K. performed the bioinformatic analysis. S.H.J. provided guidance throughout the project. All authors have read and approved the article.

## Supplementary information

Supplementary Figs. 1–3 and Tables 1-5 are provided in the Supplementary materials.

