## Supplementary Materials for "Massively parallel screening of TIR-derived peptides reveals vast TLR-targeting immunomodulatory peptides"

**This PDF file includes:**

Figures S1 to S3

Tables S1 to S5

Captions for Data S1 to S4

**Other Supplementary Materials for this manuscript include the following:**

Data S1 to S4

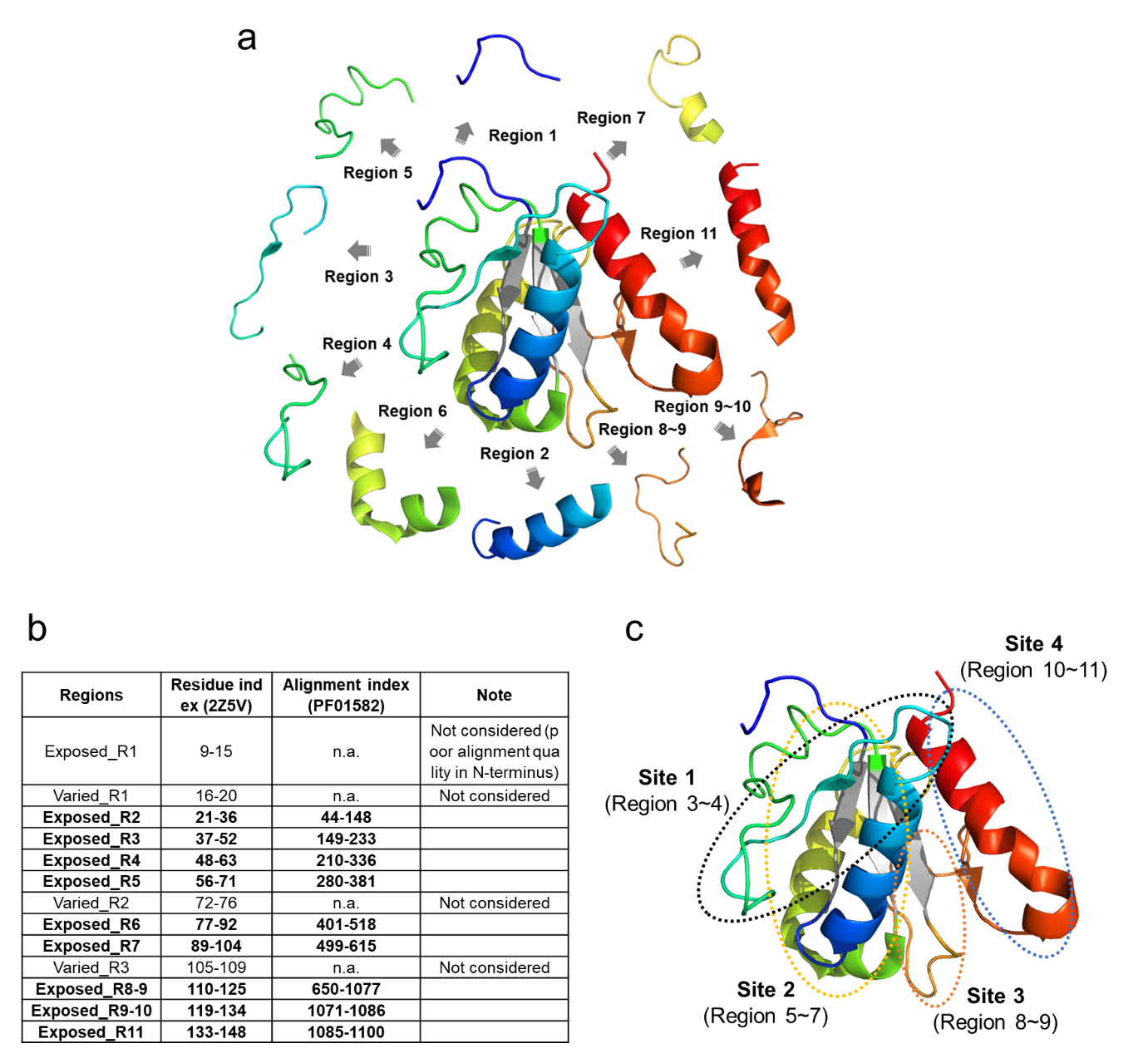

Figure. S1. Structural elements of MyD88 TIR domain. (a) Surface-exposed regions 1 to 11 of MyD88 (PDB ID 2Z5V) are shown in different colors. The hydrophobic core formed by parallel β-sheets (βA, βC and βD), which is excluded in the library design, is colored in gray. (b) The extracted regions of TIRs are summarized. (c) Four sites of TIR interfaces, which mediate the interactions between TIR domains, are marked.

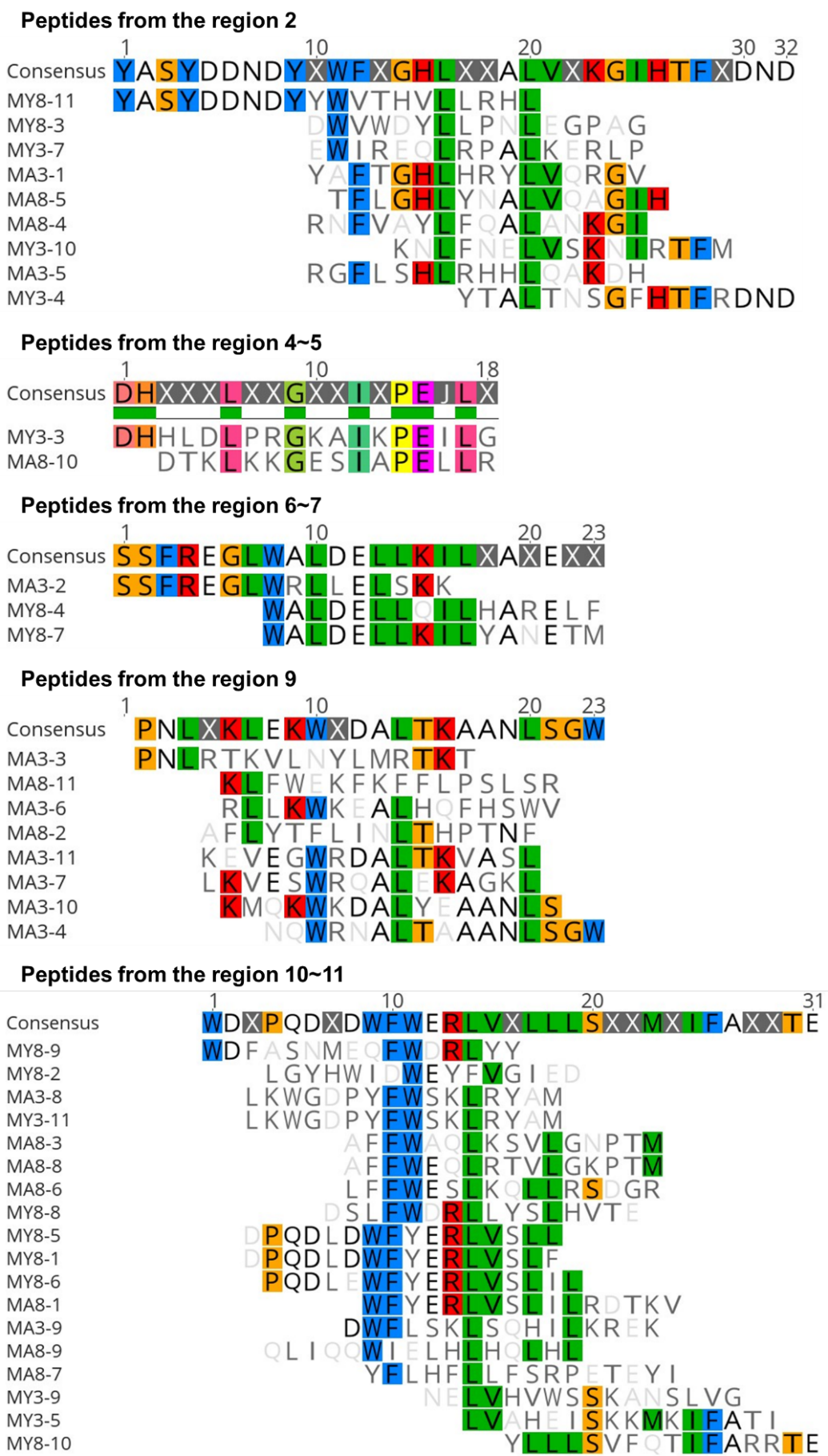

**Figure. S2.** **Sequence alignment of the selected TDIPs.** The selected peptides were aligned according to the structural regions from which the peptides originated.

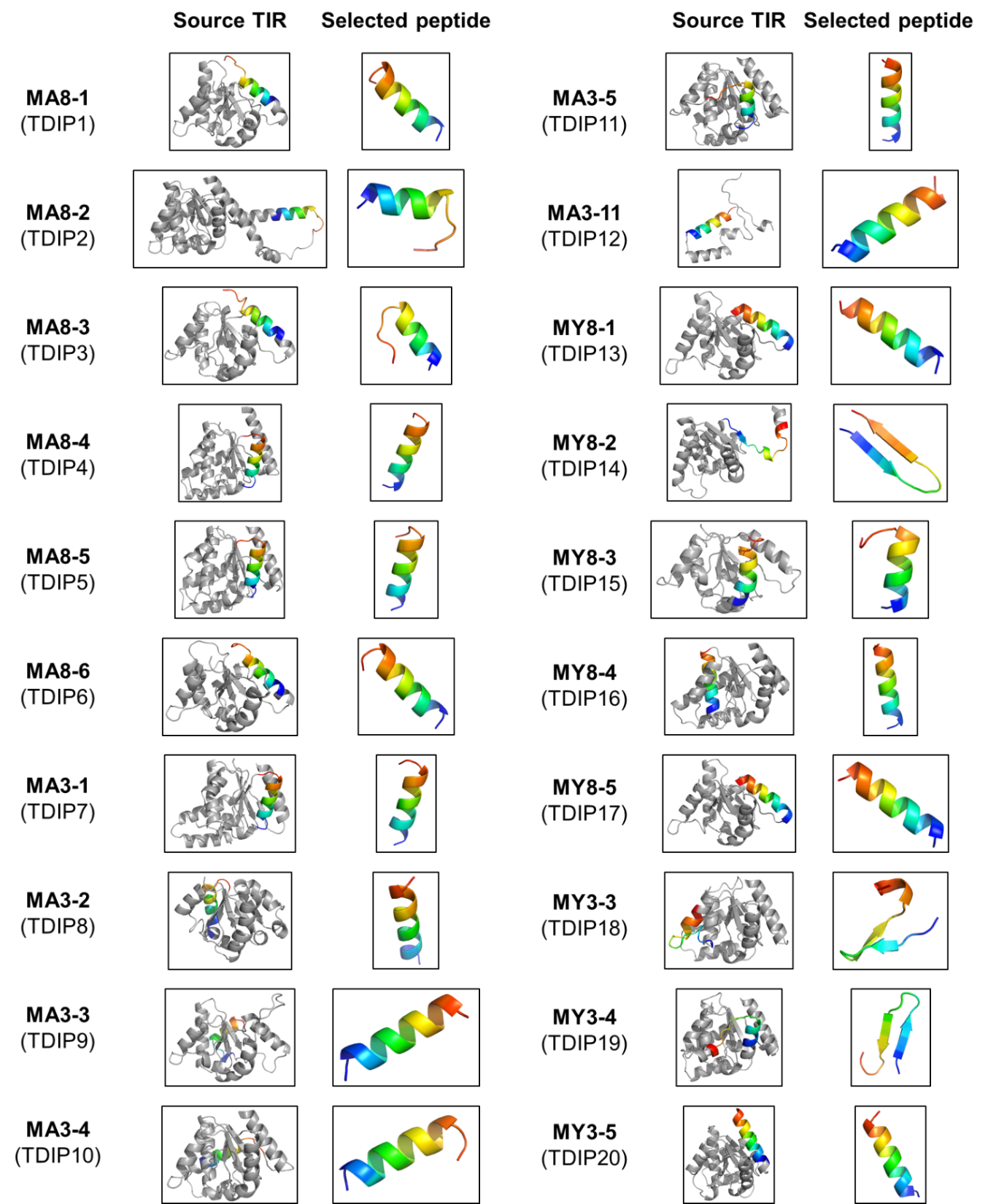

**Figure. S3.** **Structural description of the selected synthetic peptides.** The selected peptides are rainbow-colored in each AlphaFold-predicted or swiss model structure of the source TIR domains from Uniprot DB. The three-dimensional structures of the selected peptides were predicted by using PEP-FOLD4.

**Table S1. Hand-curated domains included in the T-Surf library.**

| **Uniprot ID** | **Uniprot Entry Name** | **Protein name** | **Gene Names** | **Organism** |
| --- | --- | --- | --- | --- |
| P14778 | IL1R1_HUMAN | Interleukin-1 receptor type 1 | IL1R1 | Homo sapiens (Human) |
| Q15399 | TLR1_HUMAN | Toll-like receptor 1 | TLR1 | Homo sapiens (Human) |
| O60603 | TLR2_HUMAN | Toll-like receptor 2 | TLR2 | Homo sapiens (Human) |
| O15455 | TLR3_HUMAN | Toll-like receptor 3 | TLR3 | Homo sapiens (Human) |
| O00206 | TLR4_HUMAN | Toll-like receptor 4 | TLR4 | Homo sapiens (Human) |
| O60602 | TLR5_HUMAN | Toll-like receptor 5 | TLR5 | Homo sapiens (Human) |
| Q9Y2C9 | TLR6_HUMAN | Toll-like receptor 6 | TLR6 | Homo sapiens (Human) |
| Q9NYK1 | TLR7_HUMAN | Toll-like receptor 7 | TLR7 | Homo sapiens (Human) |
| Q9NR97 | TLR8_HUMAN | Toll-like receptor 8 | TLR8 | Homo sapiens (Human) |
| Q9NR96 | TLR9_HUMAN | Toll-like receptor 9 | TLR9 | Homo sapiens (Human) |
| Q99836 | MYD88_HUMAN | Myeloid differentiation primary response protein MyD88 | MYD88 | Homo sapiens (Human) |
| P58753 | TIRAP_HUMAN | Toll/interleukin-1 receptor domain-containing adapter protein | TIRAP MAL | Homo sapiens (Human) |
| Q8IUC6 | TCAM1_HUMAN | TIR domain-containing adapter molecule 1 | TICAM1 (TRIF) | Homo sapiens (Human) |
| Q86XR7 | TCAM2_HUMAN | TIR domain-containing adapter molecule 2 | TICAM2 (TRAM) | Homo sapiens (Human) |
| Q9Y4K3 | TRAF6_HUMAN | TNF receptor-associated factor 6 | TRAF6 | Homo sapiens (Human) |
| A0A0H2V8B5 | TCPC_ECOL6 | NAD(+) hydrolase TcpC | TcpC | Escherichia coli O6:H1 (strain CFT073 / ATCC 700928 / UPEC) |
| C0RGW8 | BTPA_BRUMB | Probable 2' cyclic ADP-D-ribose synthase TcpB | TcpB | Brucella melitensis biotype 2 (strain ATCC 23457) |
| Q01220 | A52_VACCW | Protein A52 | A52R | Vaccinia virus (strain Western Reserve) (VACV) |
| P26672 | A46_VACCW | Protein OPG176 | A46R | Vaccinia virus (strain Western Reserve) (VACV) |

**Table S2. Summary of the designed T-Surf library.**

| **Designed T-Surf library** | |
| --- | --- |
| Total number of TIR domain family (Pfam PF01582) | 13,644 |
| Number of domains containing at least one peptide in the designed library | 13,603 (99.7%) |
| Total number of amino acids of TIR domain family (Pfam PF01582) | 2,164,918 |
| Number of amino acids that T-Surf library covers | 1,529,164 (70.6%) |

**Table S3. Statistics of the T-Surf library.** The number of designed peptides in the P8 and P3 libraries are summarized. The coverage and the total diversity of the libraries are given at the peptide level.

| **NGS analysis** | | | | | |
| --- | --- | --- | --- | --- | --- |
| **Library** | **Designed peptides** | **Validated peptides** | **Coverage** | **Peptide variants ^a^** | **Total diversity** |
| **T-Surf P8** | 190,945 | 190,419 | 99.7% | 997,373 | 1,187,792 |
| **T-Surf P3** | 190,945 | 104,572 | 54.8% | 603,588 | 708,160 |

a) Non-designed peptides with mutations that were incorporated during the oligo synthesis or cloning steps.

**Table S4. The redundant peptides in the T-Surf P8 library.**

| **T-Surf P8** (Total NGS count: 25,464,130) | | | |
| --- | --- | --- | --- |
| **Peptide sequence** | **Peptide ID** | **NGS count** | **Population** |
| ANAIKHANITTFFDDD | A0A2J6JQE8_LACSA/23-207_[21-36] | 6115 | 0.0240% |
| KQKHGSIRWKEDSAEK | A0A402E7G2_9SAUR/761-938_[122-137] | 6100 | 0.0240% |
| ELVKIMEAKEKEIGHI | A0A3Q7EFL2_SOLLC/18-194_A0A3Q7EFL2.1_[7] | 5945 | 0.0233% |
| RNGFAGHLYKALARRK | V7C5I8_PHAVU/11-182_[12-27] | 5861 | 0.0230% |
| SRKYLASKWRDFELNM | K1QVT6_CRAGI/4-147_K1QVT6.1_[6] | 5835 | 0.0229% |
| DVFVSFRGEDIRRRLL | A0A151SCE5_CAJCA/10-135_[2-17] | 5723 | 0.0225% |
| DVFLNFRGIDLRSGFL | A0A371HPB5_MUCPR/11-183_[2-17] | 5671 | 0.0223% |
| SEFYDVDPSEVIERKR | A0A3N7G1G5_POPTR/21-210_[100-115] | 5668 | 0.0223% |
| NGNTSYHLALHYRDIP | A0A194RA63_PAPMA/1026-1131_A0A194RA63.1_[3] | 5526 | 0.0217% |
| QVQNRWSRALSHVSNI | D7KZ23_ARALL/16-193_[122-137] | 5300 | 0.0208% |
| RGIATFQDGQLSRGIA | A0A2P6PS83_ROSCH/23-202_[24-39] | 5262 | 0.0207% |
| LAELKSSYLPVTNYLV | A0A059CQT6_EUCGR/93-197_[87-102] | 5249 | 0.0206% |
| GEIEGGHGQRYTGRFS | A0A2T7NI39_POMCA/704-863_[26-41] | 5168 | 0.0203% |
| EELMPEANRNVYGDND | W4XXW4_STRPU/733-878_[20-35] | 5086 | 0.0200% |
| ANFTGMVFKDGYESKF | A0A2J6LLJ3_LACSA/14-203_[148-163] | 5047 | 0.0198% |
| NELAMLAKLKSSLDRR | D7KPJ9_ARALL/175-350_D7KPJ9.1_[7] | 5037 | 0.0198% |
| HFKLAVHFRDFLAGIP | R7UWB2_CAPTE/8-141_[28-43] | 4764 | 0.0187% |
| VQKWRDALKDIANLSG | A0A4P1RJZ4_LUPAN/14-193_[127-142] | 4733 | 0.0186% |
| LWRDSLREVANLSGLD | A0A0R0J0J4_SOYBN/16-148_[114-129] | 4571 | 0.0180% |
| GPTDNLDPELKTYLSM | A0A195CSW9_9HYME/844-999_[100-115] | 4568 | 0.0179% |
| LQDGYESQFIQTIVKE | B9RVC7_RICCO/19-194_[145-160] | 4528 | 0.0178% |
| DELVKMNKLADLGKLR | R0H4S0_9BRAS/197-347_R0H4S0.1_[7] | 4389 | 0.0172% |
| SFRGNDVRDGFLGKLY | G7JLU8_MEDTR/9-196_[6-21] | 4381 | 0.0172% |
| GWELKNTANGGILSKL | A0A2J6JYH2_LACSA/14-208_[137-152] | 4363 | 0.0171% |
| RDEPKISKGKSISGEL | A0A251N8H2_PRUPE/18-195_[33-48] | 4301 | 0.0169% |
| DRELPNGEEISPRLYK | A0A2U1MQH6_ARTAN/11-134_A0A2U1MQH6.1_[4] | 4239 | 0.0166% |
| EHFRALGMIEKVQRWR | A0A1U8ALK0_NELNU/16-194_[118-133] | 4231 | 0.0166% |
| ALASYAGWDVRNKPEF | A0A371EM70_MUCPR/47-226_[136-151] | 4222 | 0.0166% |
| EEVSKKINRSPLHVAN | A0A1S3UVH7_VIGRR/18-195_[161-176] | 4211 | 0.0165% |
| GTDINPKLLRAIDQSM | A0A498JXL3_MALDO/209-382_[41-56] | 4183 | 0.0164% |
| HAFISYSYSDADWVRG | F6PY71_XENTR/645-786_[2-17] | 4157 | 0.0163% |
| YASSSWAARTLASYFN | A0A2P5BHT4_TREOI/23-165_[66-81] | 4136 | 0.0162% |
| GKHSPEGTSSVEYFVH | L8Y5P5_TUPCH/292-467_[8-23] | 3991 | 0.0157% |
| EFPSILRFITIADYTN | A0A2Y9LJK0_DELLE/163-295_[100-115] | 3940 | 0.0155% |
| STVLNESLKNRGINTF | A0A2G3AS97_CAPCH/90-165_[3-18] | 3878 | 0.0152% |
| LLSITKYPIGLESRVQ | G7L9E6_MEDTR/10-205_[168-183] | 3870 | 0.0152% |
| RLALKEVGNISGWHFH | A0A2I4ERM2_JUGRE/26-210_[131-146] | 3857 | 0.0151% |
| REDIEKVQGWRDALTK | A0A498K2Y2_MALDO/23-202_[120-135] | 3839 | 0.0151% |

Table S5. The redundant peptides in the T-Surf P3 library.

| **T-Surf P3** (Total NGS count: 11,882,223) | | | |
| --- | --- | --- | --- |
| **Peptide sequence** | **Peptide ID** | **NGS count** | **Population** |
| WVMSELIPQVEGEQGW | W5LU97_ASTMX/805-959_[14-29] | 4615 | 0.0388% |
| EDEAQEHYAAQDHQTE | H2ME49_ORYLA/793-971_[155-170] | 2921 | 0.0246% |
| NMAGWHVPVTRSKSKA | W9S3W0_9ROSA/4-183_[137-152] | 2304 | 0.0194% |
| VKRLDHGDSIPEELVK | A0A3Q7HJP3_SOLLC/14-198_A0A3Q7HJP3.1_[4] | 2295 | 0.0193% |
| SYANEDRGWVLDHLLP | A0A2A4K1U2_HELVI/595-746_[6-21] | 2138 | 0.0180% |
| YYRVRKLVKKLTYLSW | A0A3Q3GVF1_9LABR/763-917_[103-118] | 2131 | 0.0179% |
| WAFSNGHEAKFIHNIV | A0A251MW86_PRUPE/10-188_[142-157] | 2111 | 0.0178% |
| RELSKILEAMEARGGV | A0A2P6PKL6_ROSCH/19-206_A0A2P6PKL6.1_[7] | 2025 | 0.0170% |
| SENYAFSTWALDELVK | A0A445BMJ9_ARAHY/58-235_A0A445BMJ9.1_[6] | 1893 | 0.0159% |
| WREALAEVAALAGMVL | A0A251QIA6_PRUPE/17-198_[129-144] | 1844 | 0.0155% |
| EELVKIMERRRSFGQV | A0A2P5CSC9_PARAD/21-195_A0A2P5CSC9.1_[7] | 1827 | 0.0154% |
| NTDQSPPYNSRFWKSL | A0A3Q0DKW3_TARSY/78-241_[124-139] | 1812 | 0.0152% |
| FYGRDTRRGFTDHLRA | A0A2I4E9C4_JUGRE/24-205_[7-22] | 1806 | 0.0152% |

Data S1.

Final T-Surf library (related to Figure 1).

Data S2.

NGS data of 4R selection against MAL and MyD88 (related to Figure 1).

Data S3.

Top 11 peptides from each selected pool (related to Figure 1).

Data S4.

Peptide ranking after grouping (related to Figure 1).
